# Identification of the Repressive Domain of the Negative Circadian Clock Component CHRONO

**DOI:** 10.1101/754168

**Authors:** Yu Yang, Ning Li, Jiameng Qiu, Honghua Ge, Ximing Qin

## Abstract

Circadian rhythm is an endogenous, self-sustainable oscillation that participates in regulating organisms’ physiological activities. Key to this oscillation is a negative feedback by main clock components Periods and Cryptochromes that repress transcriptional activity of BMAL1/CLOCK complexes. In addition, a novel repressor, CHRONO, has been identified recently, but details of CHRONO’s function during repressing the circadian cycle remain unclear. Here we report that a domain of CHRONO mainly composed of α-helixes is critical to repression through the exploitation of protein-protein interactions according to luciferase reporter assays, co-immunoprecipitation, immunofluorescence, genome editing and structural information analysis via circular dichroism spectroscopy. This repression is fulfilled by interactions between CHRONO with a region on the BMAL1 C-terminus where Cryptochrome and CBP bind. Our results indicate that CHRONO and PER differentially function as BMALA/CLOCK-dependent repressors. We further suggest a repression model of how PER, CRY and CHRONO proteins associate with BMAL1, respectively.

## Introduction

Most organisms have evolved an endogenous circadian clock to adjust their metabolic activities to anticipate daily environmental events, such as day/night switches or temperature changes. In mammals, circadian clocks exist in almost every cell and organ. The clock consists of a transcriptional and translational feedback loop (TTFL) in which transcription factors brain and muscle arnt-like protein-1 (BMAL1) and circadian locomotor output cycles kaput (CLOCK) form hetero-dimer complexes (B/C complexes in the following text) and then activate transcription of clock controlled genes (ccgs), including genes encoding PER (PER1, and PER2, and PER3) and CRY proteins (CRY1, and CRY2) (1–3). After translation and post-translational modifications, PERs and CRYs translocate into the nucleus to repress BMAL1/CLOCK complex function (4, 5). With such repression, a full clock cycle is completed. A second loop composed of circadian components, such as *rev-erb* (6) and *ror* (7) and their coding products is involved in regulating transcription of the *arntl1* gene, which encodes the BMAL1 protein. Other clock genes, such as*csnk1d*, *csnk1e, fblx21, and fblx3*, contribute to setting the clock speed (8–11) or in entraining the clockwork (12).

Recently, a novel clock gene was discovered using ChIP-seq (and computational analysis) and subsequently analyzed in mammals. It was named ChIP-derived repressor of network oscillator (*chrono*) (13), or computationally highlighted repressor of the network oscillator (14), or circadian associated repressor of transcription (*ciart*) (15). *chrono* is under control of the clock and its translation product, CHRONO (CHR in the following text), is reported to repress transcriptional activating B/C complexes in a histone deacetylase (HDAC)-dependent manner (13). *chrono* deficient mice had longer period of the locomotor activity, similar to some other negative clock components such as *cry*, indicating it is a core circadian component (14). In summary, negative components required to complete a full clock cycle include PERs, CRYs, and CHR as well.

Negative repression is critical to the circadian clock and some means of repression have been identified. First, repressors PERs (PER1 and PER2) were found to remove B/C complexes from chromatin and then inactivate transcription (16, 17). In contrast, ChIP assays to map chromatin association by PERs indicated that PER forms large complexes on chromatin during early repression phase (18, 19). Studies revealed that PER2 and CRYs bind to different domains of the BMAL1 protein in order to function as repressors (20). The photolyase superfamily protein CRYs are more potent than PER proteins for repressing B/C complexes (21, 22, 25). Partch lab demonstrated that besides binding to BMAL1, CRY1 also associates with CLOCK as a repressor (24). During the repression phase, CRYs bind to B/C complexes at the C terminus of BMAL1 on chromatin, sequestering the binding site from transcriptional co-activators, CBP/p300 (25). Biochemical and cell-based complementation assays suggest that an α-helical domain of CRY1 is required for feedback repression and that the C-terminal domain of CRY1 is dispensable for complementation of the circadian clock in *cry1*/*cry2* double knockout cells (22). Repression of B/C complexes by CHR has been studied and CHR is reported to be similar to CRY1 with regard to circadian clock regulation, via interacting with the BMAL1 C terminus (23). Anafi et al. reported that CHR functioned at an independent region from (but adjacent to) CRY proteins on the BMAL1 C terminus (14). However, how repression during the cycle of the circadian clock happens remains unclear, for example, how CHR fulfills its repressor role is elusive. We do not know which region of CHR associates with BMAL1 or the structure of CHR. Thus, to understand how CHR represses B/C complex transcriptional activity, we purified a predicted helical domain of the CHR protein and extended helical domains. Then, we used circular dichroism spectroscopy to confirm secondary structures. We found that the helical domain of CHR is necessary to repress B/C complexes *in vitro*. Immunoprecipitation and Immunofluorescence revealed interactions between the middle area of CHR containing rich α-helices and the C terminus of BMAL1, indicating that CHR repressed B/C complexes via its middle area containing an α-helix rich region. Collectively, characterizations of the helical domain of CHR tell us more about how CHR represses B/C complex transcriptional activity, compared to the other two negative components (PERs, CRYs).

## Materials and Methods

### Strains and Constructs

Polymerase chain reaction (PCR) was used to make constructs described in this study. Optimized sequence coding for CHR-H and CHR-H3 were amplified and inserted into the V28-E4 plasmid (from Dr. Ge lab, Anhui University) between Not I and Xho I restriction sites, resulting in a maltose binding protein (MBP) tag and 6XHis tag fused to the 5’ and 3’ ends respectively. MBP-CHRH (or H3) was expressed in the *E. coil* BL21 (DE3) strain. For the Luciferase reporter assay, various versions of *chrono* as indicated in the text were cloned into pcDNA3.1(+)/Hygro (Invitrogen, UntiedStates). For immunoprecipitation, sequence of 5Myc-6His-hBMAL1 (5M6HA1) was cloned and inserted into pcDNA3.1 via BmaH I and EcoR I sites (named pcDNA3.1-5M6HA1 afterwards). And on the one hand, a series of *chrono* with cpVenus tag followed sequences were cloned and inserted into pcDNA3.1(+)/Hygro plasmid between HindIII and Xho I sites, on the other hand, we built series of *bmal1* with 5XMyc-6XHis-tag, using two EcoR I sites on the mutated pcDNA3.1

### Site-directed Mutagenesis

To make L06A/L607A double mutations in the bmal1 coding region, the whole plasmid pcDNA3.1-5M6HA1 was amplified using a high fidelity Taq polymerase. A pair of complementary primers 5’GGCAGCAATGGCTGTCATCATGAGCGCAGCAGAAGCAGATGCTGGACTGGGTGGC3’ and 5’GCCACCCAGTCCAGCATCTGCTTCTGCTGCGCTCATGATGACAGCCATTGCTGCCT3’ were used for the mutagenesis. After the amplification, Dpn Iwas used to digest the template plasmids. Then the product was transformed into the DH5α competent cells. To obtain the site-directed mutagenesis plasmids, the plasmids from the transformed *E.coli* colonies were sent to sequencing for identification.

### Secondary Structure Prediction

PredictProtein (www.predictprotein.org/), PSIPRED (http://bioinf.cs.ucl.ac.uk/psipred/), and JPred (http://www.compbio.dundee.ac.uk/jpred/) were used for CHR secondary structure prediction and both offered similar results.

### Nuclear Location Signal Prediction

NucPred (https://nucpred.bioinfo.se/nucpred/), PredictNLS (https://rostlab.org/owiki/index.php/PredictNLS),and PSORTII(https://psort.hgc.jp/form2.html), were used for CHR nuclear location signal prediction.

### Codon Optimization

Codon usage bias is a general characteristic for all species. The human chrono cDNA had a low yield using prokaryotic expression system. We used OPTIMIZER (http://genomes.urv.cat/OPTIMIZER/) and JCat (http://www.jcat.de) to obtain optimized sequences for chrono to express in E. coli, and the full optimized sequence was ordered from General Biosystems, Inc. Anhui Province, China.

### Protein Expression and Purification

BL21(DE3) E. coli cells that carry constructs for expression of MBP-CHRH/CHRH3 fusion protein were grown at 37 °C, 220 rpm to reach OD values up to 0.6-0.8. Then 0.2 mM IPTG was added to induce expression at 16 °C, 150 rpm, for 20-24 h. Cell pellets were lysed in 50 mM Tris-HCl (pH 8.0), and 200 mM NaCl. Ni-NTA Sefinose™ Resin (#C600033, BBI, Shanghai, China) was used for first His tag affinity purification. Then, Hiload 16/600 superdex (200pg) was used as gel filtration purification.

### Circular Dichroism Spectroscopy

To verify rich α-helix occupation in the secondary structure of CHR-H, we compared secondary structures of MBP-His_6_, MBP-CHRH-His_6_, and MBP-CHRH3-His_6_ with circular dichroism spectroscopy (Biologic/MOS-500). Protein buffer was changed to 50 mM sodium phosphate (pH 8.0), and then the protein sample was diluted to 0.1 mg/ml and clarified using by centrifugation (12,000 rpm, 4 °C, 10 min) before measurement. Each CD spectrum was collected at room temperature, with a scan rate of 1nm/s; scanning increment, 0.5nm; spectral bandwidth, 1.0nm; from 190-260 nm for each measurement. The spectra represent the average of 3 scans with the bassline subtracted from analogous conditions.

### Luciferase Reporter Assay

HEK-293T cells were grown in DMEM (SH30022, Hyclone/GE, United States) supplemented with 10% fetal bovine serum (P30-3302, PAN-Biotech, Germany) plus 100 IU/ml penicillin and 0.1 mg/ml streptomycin (SV30010, Hyclone/GE, United States) at 37°C, 5% CO_2_. Approximately 3 x 10^5^ cells were seeded in each well of a 6-well plate 1 day before transfection. When cells reached 70%∼90% confluency, 100 ng of the firefly luciferase reporter pP_PK2_::Fluc, 40ng of the renilla luciferase reporter pP_CMV_::Rluc, 250 ng of a hamster Bmal1 construct and 250 ng of a mouse Clock construct and 500 ng of mouse Per2 (or human CHR, CHR-N,CHRNH, CHR-H, CHRH2, CHRH3,CHRH4,CHRHC and CHR-C) were co-transfected into HEK 293T by using Lipo6000™ Transfection Reagent (C0526, Beyotime, Beijing, China). The constant amount of DNA (1,100 ng/well) was adjusted with pcDNA3.1(+)/Hygro vector as a carrier, and we measured single luminescence. Twenty-four hours after transfection, cells were washed three times with ice-cold PBS (SH30256.01, Hyclone/GE, United States), then processed with a Trans Detect Double-Luciferase Reporter Assay Kit (FR201, Transgen Biotech, Beijing, China), and measured by SpectraMax Paradigm Multi-Mode Microplate Reader (Molecular Devices, Silicon Valley, United States). Each construct was examined at least for three independent times.

### Western Blotting and Immunoprecipitation

Mouse antibodies against Myc-tag (M185-3L/M047-3, MBL, Nagoya, Japan), rabbit antibody against GFP-tag (D110008, BBI, Shanghai, China) and sheep antibody against Tubulin (ATN02, Cytoskeleton, Denver, United States) were subjected to Western blot according to the manufacturer’s protocol. For IP, HEK-293T cells transfected with the desired plasmids by using Attractene Transfection Reagent (301005, QIAGEN, Hilden, Germany), according to the fast-forward protocol, were lysed in IP buffer (10 mM Tris pH 7.5, 150 mM NaCl, 1 mM EDTA, 1% Triton X-100, 0.1% sodium deoxycholate, 1 mM PMSF and Roche complete EDTA free protease inhibitor cocktail). After quantification by using BCA Protein Assay Kit (P0012, Beyotime, Beijing, China), lysates were precleared with Protein G Agarose (P2009, Beyotime, Beijing, China) and then immunoprecipitated with mouse anti-Myc antibodies. After washing five times, the precipitates were resuspended in the 5XSDS loading dye, boiled for 5min, and run on a 12.5% SDS-PAGE gel followed by Western blot analysis. Immunoreactive bands were detected by Typhoon laser scanners (FLA-9500, GE, United States).

### Immunofluorescence and Localization

For IF, Hela cells were grown in DMEM supplemented with 10% FBS plus 100 IU/ml penicillin and 0.1mg/ml streptomycin at 37°C, 5% CO_2_. A sterile cover glass was placed into each well of a 6-well plate, and then 2 x 10^5^ cells per well were seeded 1 day before transfection. When cells reached 70%∼90% confluency, they were transfected with Lipo6000™ Transfection Reagent with various combinations of the following constructs (all in pcDNA3.1 (+)/Hygro background): 1,250 ng CHR-cpVenus or CHRH3-cpVenus and 1,250 ng 5XMyc-6XHis-hBMAL1, 5XMyc-6XHis-hBMAL1-C or 5XMyc-6XHis-hBMAL1-Cs. Twenty-four hours after transfection, cells were washed three times with ice-cold PBS and fixed in 4% paraformaldehyde for 30 min at room temperature. Then cells were permeabilized and blocked with 0.3% TritonX-100 and 5%FBS in PBS for 1h at room temperature follow by sequential washing with PBS, and were subjected for incubation using Myc-tag antibody (ab32, UK, abcam, diluted in PBS containing 3%BSA) for 1h at room temperature. By following three 5 min PBS washes, cells were immune-stained by Cy3-conjugated antibody (D110172, BBI, Shanghai, China, diluted in PBS containing 1%BSA/0.3%TritonX-100) for 30 min at room temperature. After last three times PBS wash, cells were embedded in Mounting Medium, Antifading (with DAPI, S2110, Solarbio). Analyses and documentation were done with immunofluorescence microscopy (TCS SP5, Leica, Germany).For localization assessment, Hela and NIH 3T3 cells were cultured in the same media, and a sterile cover glass was placed into each well of a 6-well plate, and then 2 x 10^5^ cells per well were seeded 1 day before transfection. When cells reached 70%∼90% confluency, they were transfected with Lipo6000™ Transfection Reagent with series of cpVenus fused truncations of CHR (all in pcDNA3.1 (+)/Hygro background): 2500 ng CHR-cpVenus/cpVenus-CHR/CHRN-cpVenus/CHRH-cpVenus/CHRC-cpVenus or EGFP. Twenty-four hours after transfection, cells were washed three times with ice-cold PBS and fixed in 4% paraformaldehyde for 30 min at room temperature. After three times PBS wash, cells were embedded in Mounting Medium, Anti fading and documented by immunofluorescence microscopy.

### *chrono* knock out in U2-OS cells

*chrono^-/-^bmal1^Luc^*cell line was generated using the CRISPR/Cas 9 system. Oligonucleotides specific for the target sites of *chrono* gene loci were designed using the Optimized CRISPR Design tool (www.genomeengineering.org/). The highest quality score sgRNA (CCGCTGCAGGCATCGATCGA) targeted to the exon 1 of *chrono* (on-target locus: chr1:+150255862) was chosen to insert into the expression vector pX459 (pSpCas9{BB}-2A-Puro was a gift from Feng Zhang, Addgene plasmid # 48139). 24 hr after transfection, cells were screened with 2 μg/ml puromycin.Cell line clones were screened by the limiting dilution method.

### Bioluminescence recoding and data analysis

2X10^5^ U2-OS reporter cells were seeded in 35mm dishes the day before synchronization. On the next day, cells in each dish were synchronized with 2 h treatment of 200nM dexamethasone (dissolved in DMSO) and recorded in a LumiCycle as described previously (26). Bioluminescence data were analyzed with the LumiCycle analysis program (Actimetrics, USA) to obtain circadian parameters such as period and amplitude.

## Results

### The helical domain of CHR represses the transcriptional activity of B/C complexes

Since there were neither homologous proteins nor similar protein structures being reported, assessment of CHR amino acid sequences was used to predict its secondary structure. PredictProtein (27) and PSIPRED (28) were used to predict that CHR contains an α-helix rich region in the middle (Fig. 1A) and both ends of CHR are flexible regions. Thus, CHR was separated into three domains: the N-terminus (CHRN, 1-110), the helical (CHRH, 111-196), and the C-terminus domains (CHRC, 197-385).

**Figure 1.**
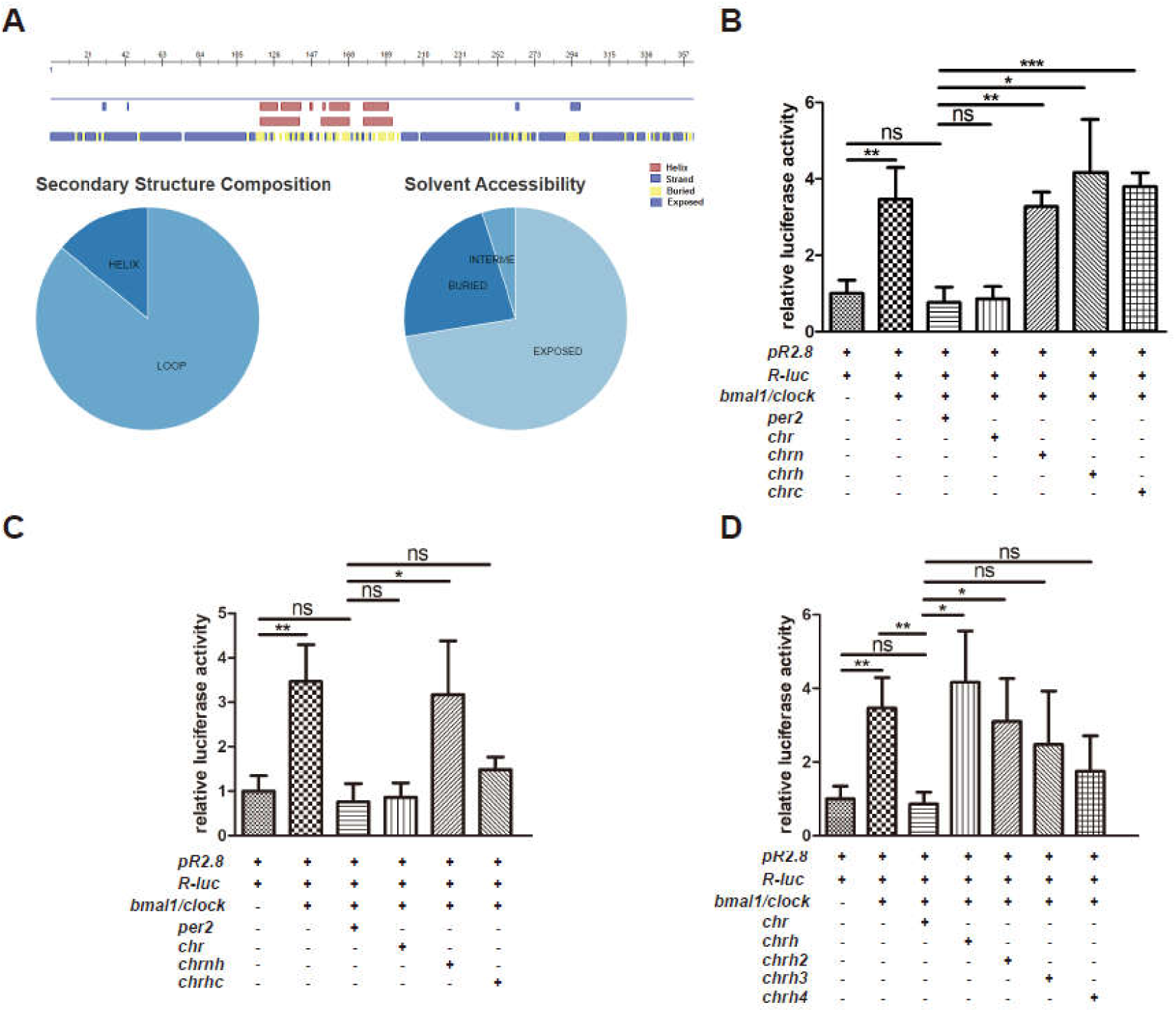
CHR repression domain that contains an α-helix rich region. **(A)** Predicted secondary structures show an α-helix rich region in the middle of CHR, with flexible regions in the N- and C-termini. Pie charts show that CHR containing a large proportion of flexible composition and a lot of regions are exposed. **(B)**Based on predicted secondary structures, CHR was divided into 3 separate parts: CHRN, CHRH, and CHRC. In the classical luciferase reporter assay which uses B/C complexes to activate an E-box containing promoter (pR2.8 in this experiment), each individual part of CHR lost the repressive activity of CHR. **(C)**However, the combination of the predicted helix region plus the C-terminus is able to function as a repressor, rather than the combination of the helix region plus the N-terminus.**(D)** Luciferase assaysindicate that CHRH plusextended regions at its C-terminus(CHRH3 and CHRH4) could repress the transcription activation by B/C complexes. Since CHRH3 is 2 amino acids longer than CHRH2 that did not perform the repressive function, this indicates that the helix containing H3 is the minimal domain that functions as a repressor to B/C complexes. At least three independent experimental repeats were done for each luciferase reporter assay. Graphpad Prism 5 was used to generate graphs/plots and perform statistical analysis (2-tailed unpaired t-test). *, *p<*0.05; **, *p<*0.01.***, *p<*0.001.

CHR was identified as a circadian clock component that functions as a transcriptional repressor of B/C complexes (13, 14). However, details of the binding regions of CHR to B/C complexes, in other words the functional domains of CHR, are unknown. Then, we searched the area of CHR that functions as an inhibitor to the transcription activator B/C complexes using the luciferase reporter assay. Based on the prediction, luciferase reporter assays were used to test domain effects including CHRN, CHRH and CHRC with respect to transcription activation by the B/C complexes. In this mammalian reporter system, both mPeriod-2 (mPer2) and full-length CHR as positive controls repressed the B/C complexes using a reporter pP_PK2_::Fluc (pR2.8) that contains E-box (CACGTG) elements in the promoter region and a reporter pP_CMV_::Rluc(R-luc) as the internal reference (29). However, these separate domains were unable to repress transcription activation by the B/C complexes individually (Fig. 1B), which indicates the repressive domain is larger than any individual part. Therefore, we performed the assays using the combinations of individual parts (CHRNH and CHRHC represent the combinations of CHRN, CHRH and CHRC) to find the inhibition part of CHR. CHRHC has been regarded as the functional area via showing repression as strong as the PER2 protein on the B/C complexes (Fig. 1C).Furthermore, a series of the extended constructs, CHRH2 (111∼262), CHRH3 (111∼264) and CHRH4 (111∼298), were made on the basis of the CHRH and CHRHC. Then, CHRH2, CHRH3 and CHRH4 were applied in the luciferase reporter assay. As the data indicate, CHRH3 and CHRH4 prevented the activation by the B/C complexes, while CHRH2 was not able to repress the activation (Fig. 1D).According to these results, the functional domain of CHR protein is in the middle of CHR whose significant part is predicted to be an α-helix rich region.

### Secondary structure of MBP-CHRH/CHRH3 measured by CD spectroscopy

Since we constructed those truncations on the basis of the secondary structure prediction, we further carried out experiments to confirm our prediction. Thus, we first attempted to acquire its tertiary structures through protein crystallization. We tried different tags (GST, 6XHis or MBP) and series of truncations to purify CHR proteins. Eventually, we chose MBP as a tag to purify truncated CHR proteins. However, we were not successful to get information of CHR’s tertiary structure through crystallization by now. Based on our luciferase reporter assays, we decided to validate the predicted secondary structure by means of CD spectroscopy. In view of yield and purity, MBP tagged CHRH and CHRH3 were chosen to carry out the CD experiments, MBP-CHRH yielded ∼ 40 mg per liter culture, while MBP-CHRH3 merely had 2 mg per liter which may be due to the increase of flexible region (Figs. 2A&2B). According to the prediction, CHRH is α-helix rich, while the component of CHRH3’s elongated part is mainly flexible (Fig. 2C).We supposed MBP-CHRH’s ratio of absorption peak at 222nm and 208nm, which characterizes the percentage of helicity (30), should be larger than that of MBP-CHRH3. Based on the result of measurement, MBP-CHRH (ratio: 1.2506) exhibited higher percentage of helicity than MBP-CHRH3 (ratio: 1.0862) (Fig.2D), and MBP-CHRH fusion did not change the helicity percentage of MBP (ratio: 1.2849) as MBP-CHRH3 did. Consistent with secondary structure prediction, CHRH is an α-helix rich region.

**Figure 2.**
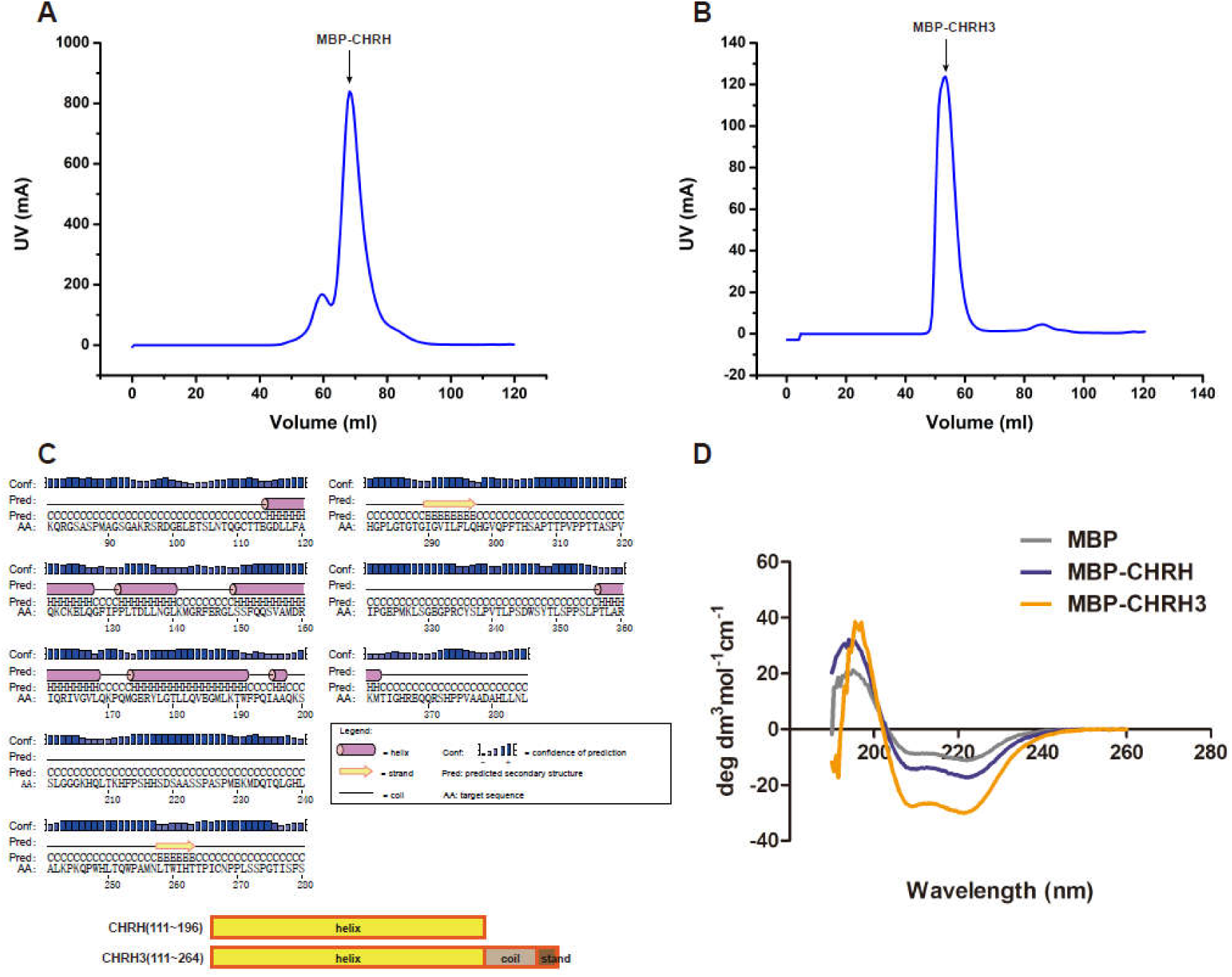
CD spectra of MBP, MBP-CHRH and MBP-CHRH3. MBP-CHRH**(A)** and MBP-CHRH3 **(B)**purified by gel filtration chromatography. Yield of MBP-CHRH was extremely higher than MBP-CHRH3’s. We gathered proteins at the peak area for subsequent experiments.**(C)**According to the result of secondary structure prediction from PSIPRED, CHRH, from 111∼196, is mainly composed of α-helix motifs; CHRH3,from 111∼264,has extra coils and β-strand motifs in comparison to CHRH. **(D)** Comparison of the CD spectra between MBP, MBP-CHRH and MBP-CHRH3 proteins. MBP and MBP-CHRH have similar contents of helixes, while MBP-CHRH3 has less helical contents, based on these CD spectra.

### The helix-rich domain is necessary for CHR/BMAL1 combining

In light of above luciferase reporter assays, we sought to confirm the interaction between CHR and BMAL1 using co-immunoprecipitation(co-IP). HEK-293T cells were co-transfected with BMAL1 which is fused to a 5Myc-6His tag (5M6HA1, for short of 5Myc-6His-BMAL1) on the N-terminus and CHR which is fused to cpVenus on the C-terminus (CHRV, for short of CHR-cpVenus). The C-terminus fusion with a cpVenus tag did interfere with its cellular localization (Fig. 6A). In consistent with previous reports that CHR interacts with BMAL1 (13–15), CHRV could be co-immunoprecipitated from extracts of HEK-293T cells together with BMAL1 proteins using an anti-Myc antibody (Fig. 3A). Subsequently, we tested the possible direct interactions between BMAL1 and series of CHR constructs as shown in Figure 1. In line with previous luciferase reporter assays, the co-IP results indicated that CHRN, CHRH or CHRC alone fused to cpVenus (CHRNV, CHRHV or CHRCV) was not able to interact with 5M6HA1 (Fig. 3B). Again, in agreement with the reporter assays, the detection of CHRHC-cpVenus (CHRHCV) in the co-IP products using anti-Myc antibodies indicated that CHRHC carries the repressive function and the helical domain of CHR is necessary for BMAL1/CHR combining (Fig. 3C). Moreover, in the same system, shorter CHRHC truncated variants (CHRH3 and CHRH4) could be immunoprecipitated by anti-Myc antibodies from extracts of HEK-293T cells co-transfected with 5M6HA1 (Fig. 3D&3E), while CHRH2, merely two amino acids shorter than CHRH3, could not be detected (Fig. 3F). These results signify that the middle region of CHR (111∼264) is the minimum area for BMAL1/CHR binding. In summary, it is vital for CHR to perform as a transcriptional repressor that its middle area containing a highly helical domain combining to BMAL1 directly.

**Figure 3.**
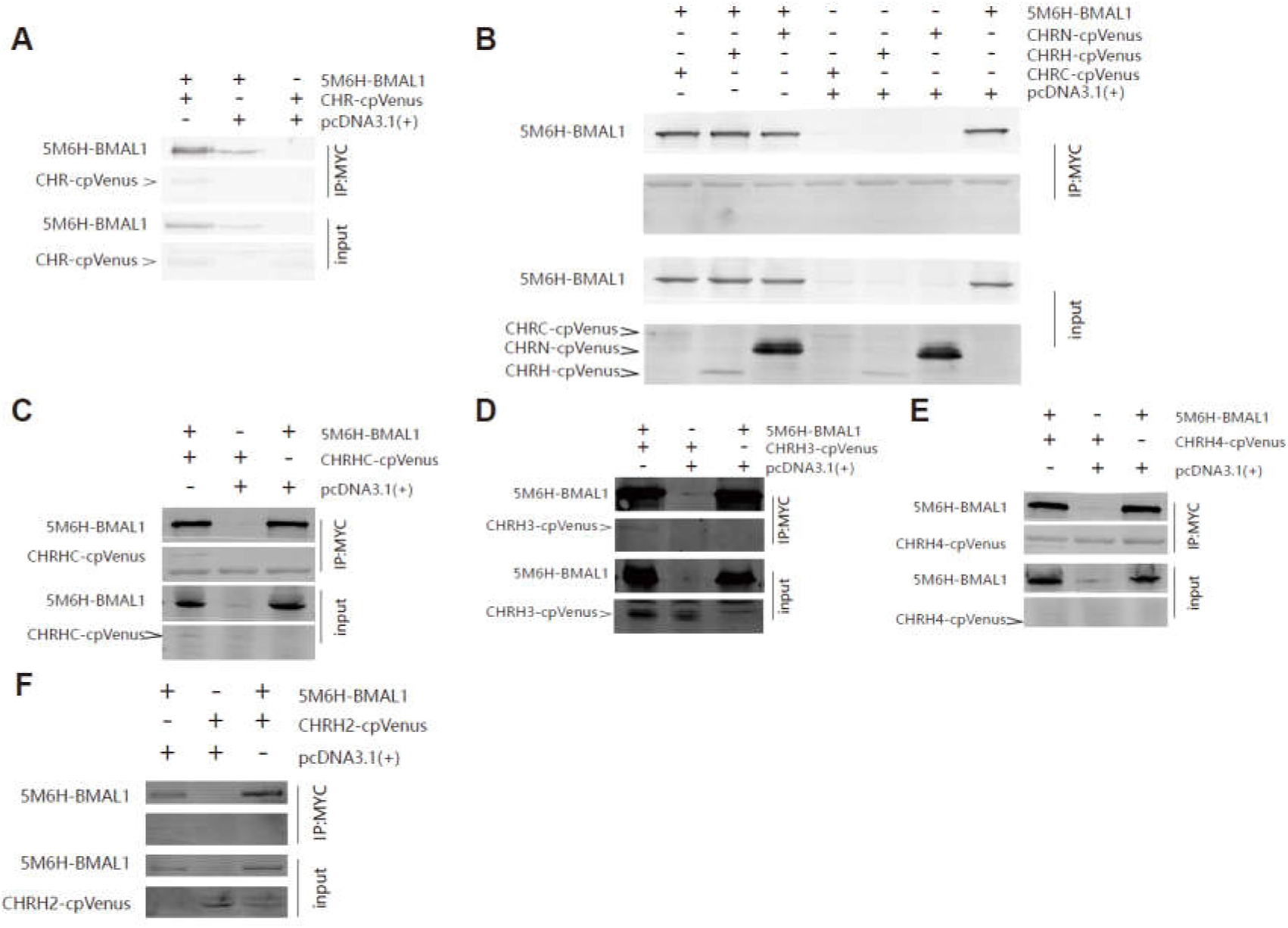
Co-IP assays indicate thatthehelix-rich domain is necessary for CHR/BMAL1 interaction. **(A)** Full-length CHR fused with cpVenus could be immunoprecipitated by full-length BMAL1 fused with a 5Myc-6His-tag when both constructs were co-transfected in HEK 293T cells. The cellular extracts were immunoprecipitated with anti-Mycantibodies and detected by using anti-GFP antibodies. **(B)** CHRN, CHRH and CHRCwere not able to be immunoprecipitated by full-length BMAL1 when they were individually co-transfected with BMAL1 in HEK 293T cells. Consistent with the luciferase reporter assays, CHRHC**(C)**and its shortened truncations, CHRH3**(D)** and CHRH4**(E)**, were immunoprecipitated by 5M6H-BMAL1when they were individually co-transfected with BMAL1 in HEK 293T cells.**(F)** CHRH2,with two amino acids cut at CHRH3’s C-terminus, failed to be immunoprecipitated by 5M6H-BMAL1.At least two independent experimental repeats were done for each co-IP experimental set.

### The C-terminus of BMAL1 that is responsible for binding with CHR

A previous study had found a region (514∼594) of BMAL1 adjacent to the CRY1 interacting terminus using sequence alignment between C-terminal domains of BMAL1 and BMAL2 (14).The full-length CHR was reported to interact with this region, causing interference with binding between B/C complexes and CRE binding protein (CBP) or p300, the transcriptional co-activator of circadian clock pathways (14). However, they did not furnish evidence that the unique region alone has an interaction with CHR by means of immunoprecipitation or other reliable interaction assays. To better understand how CHR interacts with BMAL1 to repress the activity of B/C complexes, we decided to confirm the interaction between CHR and BMAL1C. However, in our data, 5Myc-6His-hBMAL1-C (5M6HA1-C, 514∼594) failed to co-immunoprecipitate with the full-length CHR protein (Fig. 4A).In addition, to solidify our evidences, we also carried out immunofluorescence (IF) assays to test the interactions (Fig. 4H). As a transcriptional repressor, exotic expressed CHRV located to the nuclei, as detected by the fused fluorescent protein. Our IF assay using anti-Myc antibodies indicated that BMAL1-C located to the cytoplasm. The merged image implied that CHRV and 5M6HA1-Cwere not able to interact with each other (Fig. 4H), consistent with our co-IP results (Fig. 4A).Under such circumstance, we believed that there is no sufficient evidence that the right domain of BMAL1 C-terminus has been identified for BMAL1/CHR combination. Since we knew that the last 43 amino acids of BMAL1 are required for transcriptional activation (23)and CBP also combines to it (25), we hypothesized that CHR disturbed CBP’s co-activation by combining to the last 43 aa of BMAL1(583∼626)as well. To test our hypothesis, firstly, we tested whether CHR interacted with the C-terminus of BMAL1. Indeed, 5Myc-6His-hBMAL1-N (5M6HA1-N, 1∼490) was not capable of co-immunoprecipitating with CHRV together from extracts of HEK-293T cells (Fig. 4B), while 5Myc-6His-hBMAL1-C longer version (5M6HA1-Cl, 490∼626) succeeded (Fig.4C). Then, we used 5Myc-6His-hBMAL1-C shorter version (5M6HA1-Cs, 579∼626), containing the last 43 aa, to pull CHRV down, and it also succeeded in precipitating the CHRV protein (Fig. 4D). What’s more, owing to the fact that CRY1 and CBP compete for BMAL1 TAD domain binding (31, 32), and the fact that IxxLL motif in the TAD α-helix is the major binding site for both CRY1and CBP (25), we constructed L606A and L607A double mutated 5M6HA1 (mut5M6HA1), which destroyed the IxxLL motif (25), to exam whether CHR competes with CBP by combining to TAD domain directly. And the data illustrates that under the ruin of the IxxLL motif in the TAD domain, interaction between BAML1 and CHR disappeared (Fig. 4E).In other words, CHR, like CRY1,also has a competition with CBP for BMAL1 TAD binding. Since CHRH3 retains the repression function of CHR (Fig. 1D), we used 5M6HA1-Cs to pull CHRH3V down. Although CHRH3V has been precipitated, compared with the combinations between CHR and BMAL1-Cs and between CHRH3 and BMAL1, there is much weaker combination between CHR3 and BMAL1-Cs (Fig. 4F). It is probably that the functional domain may need extra parts to stabilize the combination. Therefore, we tested the combination between 5M6HA1-Cs and CHRH4V, which is capable to repress the transcriptional activity evoked by B/C complexes and be immunoprecipitated by BMAL1 as well (Figs. 1D & 3E). Then, a stronger binding, compared with CHRH3, has been found between CHRH4 and the C-terminus of BMAL1 (Fig. 4G). In terms of these data, we suggested that CHRH3 is the minimal domain to exercise as a repressor via being attached to BAML1-Cs, and the coils adjacent to its C-terminus might be regarded as an additional stabilizing part to maintain the binding. Similarly, we confirmed the combination between CHR and the C-terminus of BMAL1 via IF (Fig. 4H).Consistently, the short construct CHRH3 showed less co-localization, compared to the full-length CHR (Fig. 4H, bottom V.S. middle). Thus, the repressing domain, CHRH3, is important for interactions between CHR and B/C complexes through the TAD domain at BMAL1’s C-terminus..

**Figure 4.**
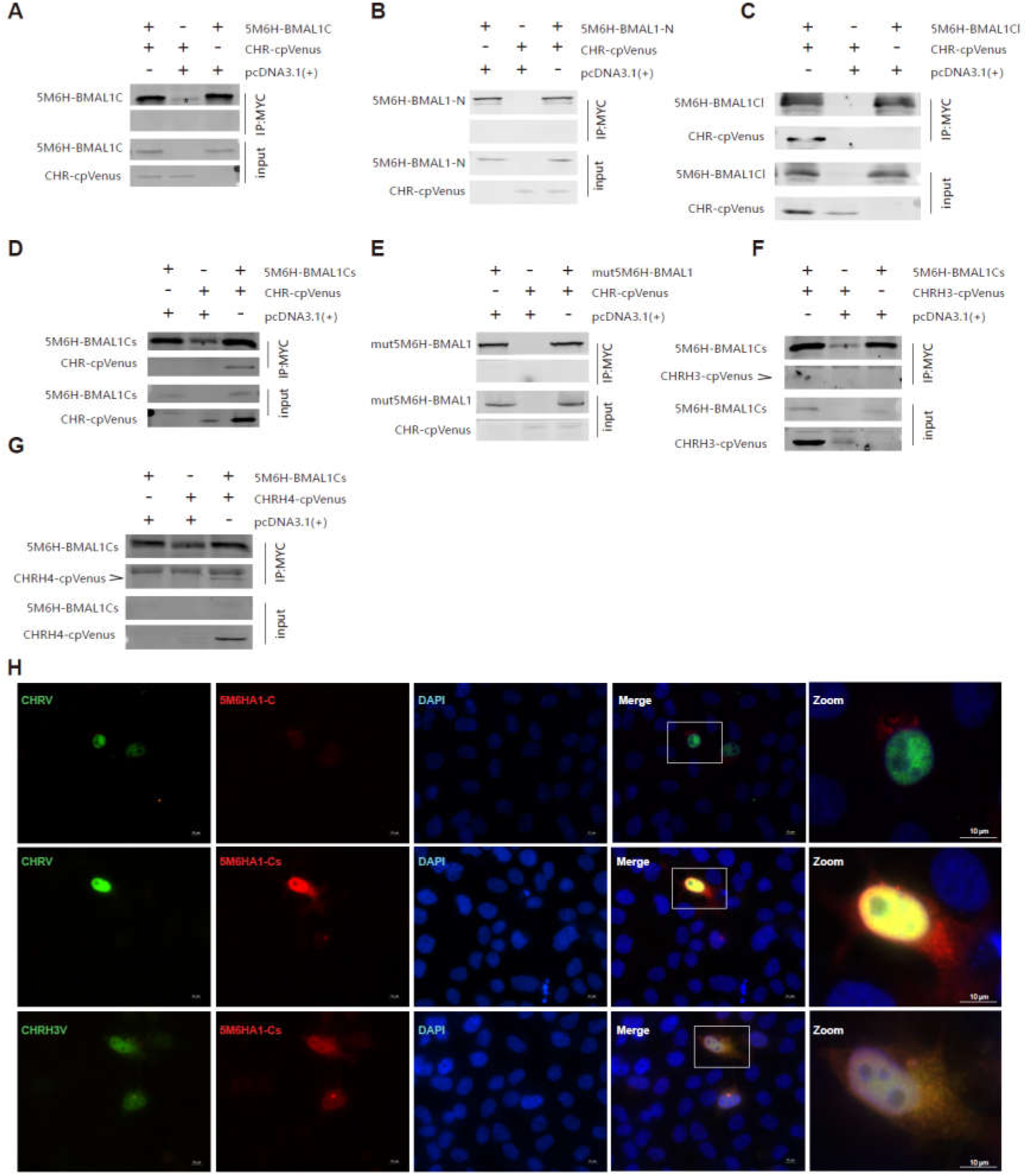
Co-IP and IF suggest that the C-terminus rather than other region of BMAL1 interacts with CHR. **(A)** The unique region of BMAL1 reported by previous study (14), from 514 to 594, could not pull CHR down in HEK 293T cell extracts. **(B)**The N-terminus of BMAL1 is not able to co-immunoprecipitated with full-length CHR either when both were co-transfected in HEK 293T cells. **(C)**However, a construct of the intact C-terminus of BMAL (from 490 to 626) named as BMAL1Clong (BMAL1Cl) is able to precipitate full-length CHR in HEK 293T cells. **(D)**BMAL1-C short (BAML1Cs, from 579 to 626),the distal C-terminal domain of BMAL1, is sufficient to immunoprecipitate CHR from HEK 293T cell extracts when both BMAL1Cs and CHR were co-transfected. **(E)**Via disruption of the IxxLL motif in the TAD domain at BMAL1’s distal C-terminus via Leucine-to-Alanine mutations at 606 and 607 sites, full-length CHR was not able to be precipitated using anti-Myc antibodies that successfully pulled down 5M6H tagged BMAL1 proteins. **(F)**BMAL1s could weakly precipitate with the minimum repressing domain of CHR (CHRH3) using anti-My cantibodies, which agrees to the luciferase reporter assays. **(G)**In comparison to CHRH3, more CHRH4 was precipitated by BMAL1Cs.**(H)**In line with these IP results, immunofluorescence (IF) confirmed the interactions between BMAL1-Cs and full-length CHR and the minimum repressor CHRH3, while CHR failed to co-localize with BMAL1-C (514∼594).Note that an unspecific band appeared when the short forms of BMAL1 C-terminus was used in the IP assays. Asterisk marks in panels A, D and F designate unspecific precipitated products using anti-myc antibodies that ran at the same position with 5M6H-BMAL1C or short version of BMAL1C.At least two independent experiments were carried out as repeats.

### CRISPR knocking-out of CHR disturbs the circadian timekeeping

As one of the core components of mammalian molecular circadian clock, BMAL1’s C-terminal TAD domain determines circadian period (25). Since CHR interacts with BMAL1’s TAD domain, we proposed that CHR also contributes to generating the 24 hr period. Previous studies have demonstrated that knocking-out *chrono* in animals could lengthen the circadian period (13–15), which agrees with our proposal. Nevertheless, it is unclear whether KO of *chrono* in cultured cells would interfere with the circadian period. Using the CRISPR/Cas9 system (33), the highest score of a 20-nt guide sequence targeting exon 1 of *chrono* gene was selected as the sgRNA, which is described in the M&M. The *chrono* gene was knocked out in U2-OS cells which harbor a luciferase reporter under the control of bmal1 promoter, generating *chrono^-/-^bmal1^Luc^* U2-OS cells. We PCR-amplified the target region from *chrono^-/-^bmal1^Luc^* genomic DNA, followed by sequencing. The results showed that both *chrono* alleles were modified, with one allele losing 4 bp and the other allele having 1 bp insertion at the Cas9 cleavage site (Fig. 5A). These frameshift mutations created premature translation stop codons, which eventually knocked out chrono in U2-OS cells (Fig. 5B). Comparing to the wild-type U2-OS cells (WT, 22.18 ± 0.068 hr), *chrono^-/-^bmal1^Luc^* cells (24.47 ± 0.453 hr) exhibited lengthened circadian period of ∼2 hours (Figs. 5C&5D). Our cell based results were in consistency with previous animal studies. Overall, these data suggested that CHR plays a role in maintaining the 24 hr periodicity via interacting with the C-terminal TAD domain of BMAL1.

**Figure 5.**
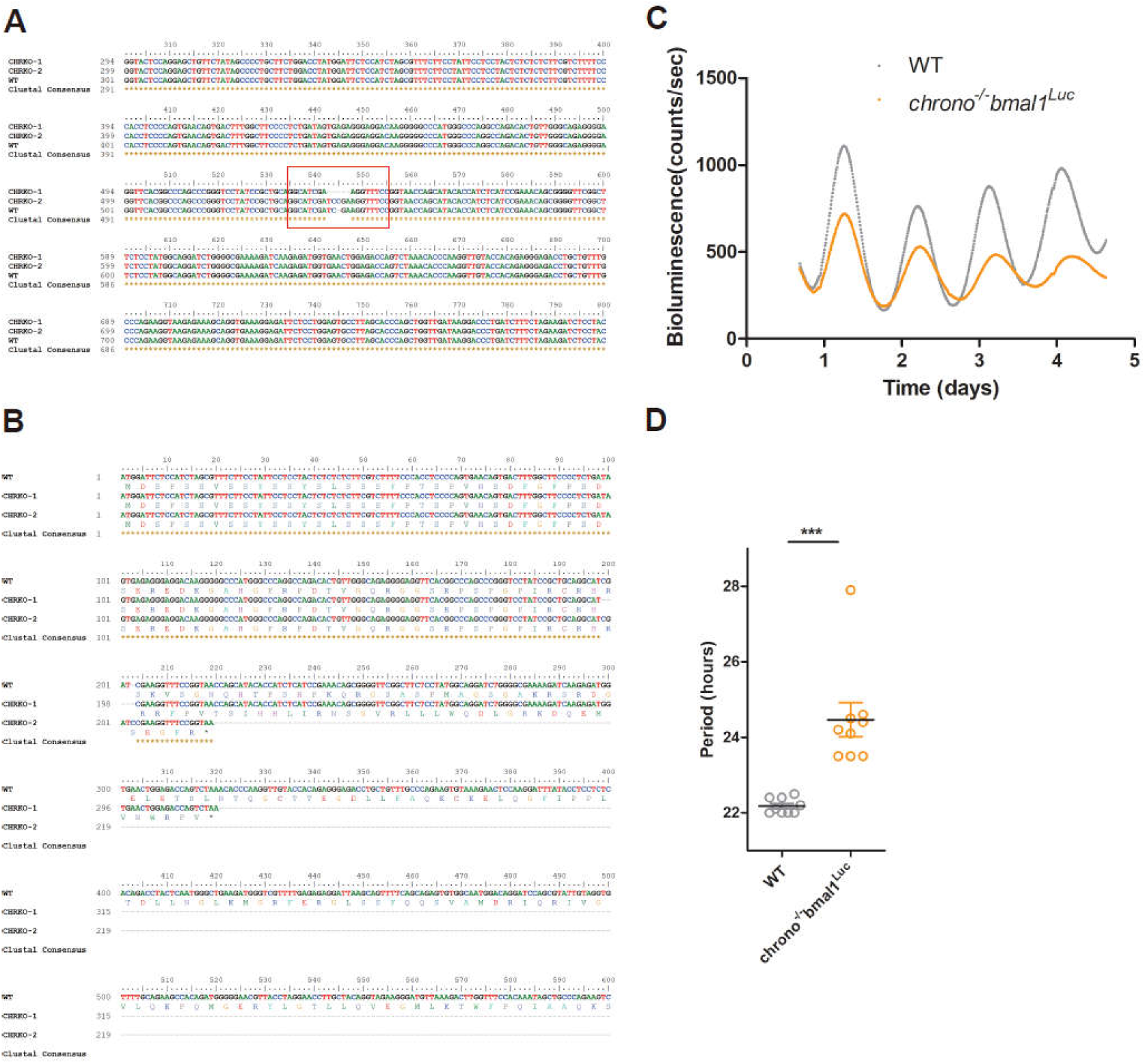
*chrono* deletion gives rise to longer period in U2-OS cell line. (A) Sequence alignment between wild-type and *chrono^-/-^bmal1^Luc^* cells showed that the knockout cell line is heterozygous with one allele deleting four bases and the other inserting one base. (B) These indels caused early termination of translation through sequence analysis. Corresponding amino acid sequences are presented underneath the nucleotide sequences. Asterisk marks indicate the stop codons caused by the indels. (C) Representatives of raw bioluminescence data from U2-OS cells expressing BMAL1:dLUC reporter are plotted for both WT and *chrono^-/-^bmal1^Luc^* cell lines. (D) Period measured by LumiCycle luminometer is plotted using nine repeats from each cell line. Graphpad Prism 5 was used to generate graphs/plots and perform statistical analysis (2-tailed unpaired t-test). ***, *p<*0.001.

### cpVenus fused to the N-terminus of CHR interferes with its nuclear localization

We have identified the domain of CHR that interacts with BMAL1 to repress the transcription activation of B/C complexes. This domain contains the predicted helix domain and a partial extended coiled region in its C-terminus. These facts make us wonder what the N-terminus of CHR functions. Thus, we constructed two plasmids that cpVenus conjugated at CHR’s C-terminus (CHRV) and the N-terminus (VCHR) separately to study the cellular localization of CHR. At first, we checked the location of those proteins in the U2-OS cell line, because CHR is a transcriptional factor located in the nuclei (13), and these two fused proteins were supposed to enter nuclei, too. However, VCHR not CHRV tended to stay at cytoplasm (Fig. 6A). On account of this phenomenon, we hypothesized that a nuclear localization signal (NLS) may lie in the N-terminus of CHR and cpVenus as a rather huge tag (26KD) may block the nuclear importin via covering the NLS or interfering with its importing partner. However, existing NLS prediction algorithms did not help us find any obvious NLS motif: although CHR is a nuclear protein, there is no specific sequence being rendered as a NLS. Perhaps there is a novel NLS in the N-terminus, so we sought to test whether CHRNV located in the nuclei. Later, we checked the expression of these cpVenus fused truncations via immunofluorescence microscopy. In Hela cell line, we found CHRNV was inclined to locate in the nuclei as much as CHRV, while CHRHV and CHRCV spread evenly in whole cell (Fig. 6B). Furthermore, we used NIH 3T3 cells to check distributions of these truncations. Compared with distributions of CHRHV and CHRCV, CHRNV has s strong accumulation in the nuclei (Fig. 6C). Noteworthy, expression of CHRNV was obviously higher than CHRHV and CHRCV’s in both cell lines, and these data are consistent with the western blot results (Fig. 3B). Perhaps, the nuclear-entrance protected CHRN from being degraded. Based on the statistics, we suggested that N-terminus of CHR leads itself to enter the nuclei, though we have not identified the exact NLS signal yet.

**Figure 6.**
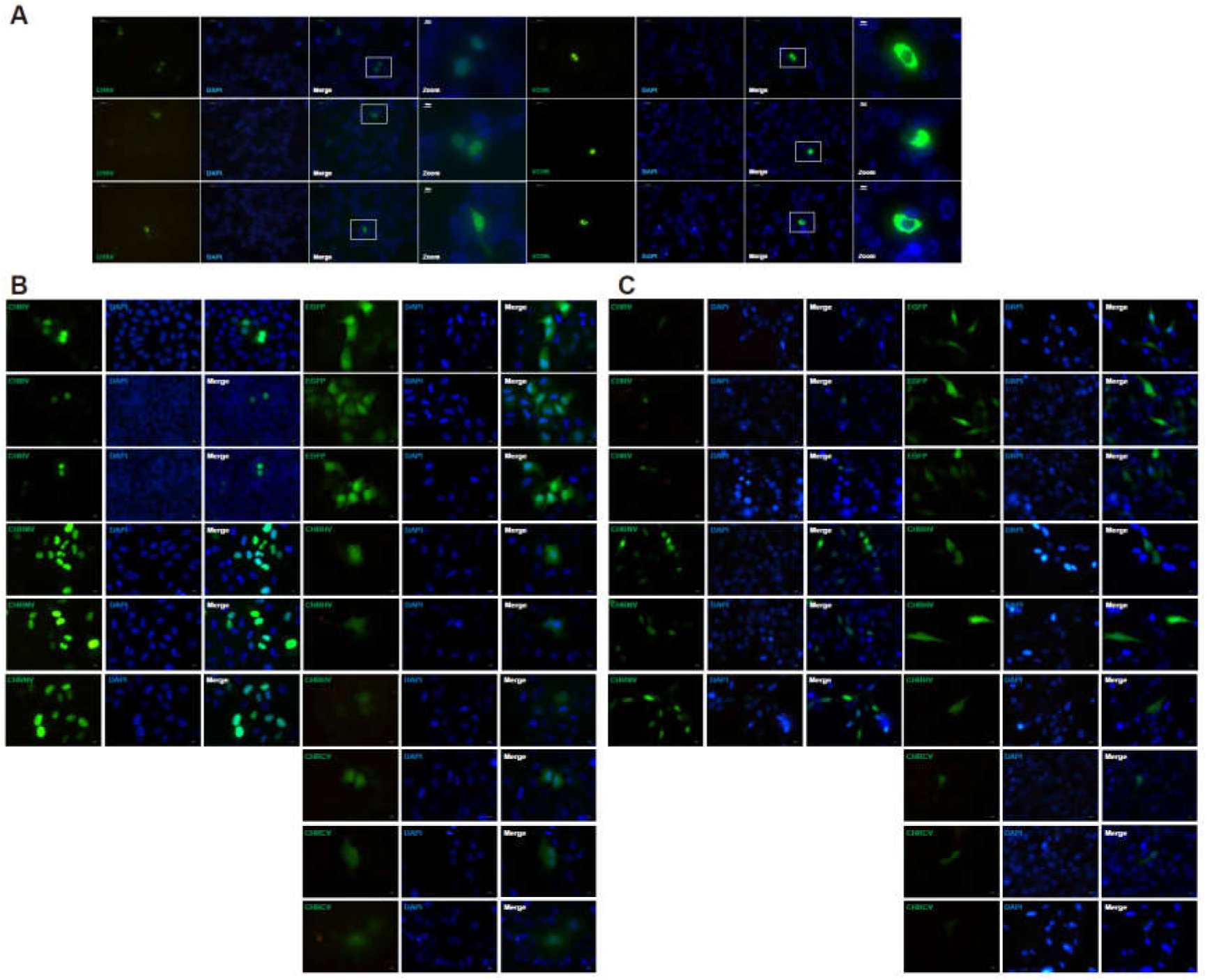
N-terminus of CHR is required for its nuclear localization. (A) Localizations of CHRV (cpVenus on the C-terminus of CHR) and VCHR (cpVenus on the N-terminus of CHR) in U2-OS cell line. CHRV mainly localized in the nuclei, while VCHR mainly stayed in the cytoplasm. (B) Different CHR constructs were expressed to study their cellular distributions in Hela cells. (C) Different CHR constructs were expressed to study their cellular distributions in NIH 3T3 cells. CHRV was used as a control to show the nuclear localization of CHR, while EGFP was used as a control to show universal localization of GFP. According to the data presented in panels B and C, the N-terminus of CHR determines the nuclear localization. Representative images are presented.

## Discussion

Analysis of “*ciart*” or “*chrono*” in the Uniprot website suggested that this gene appeared in vertebrate during evolution. No homologous gene has been reported, and no structures are described. Thus, to learn about CHR protein, we optimized codon usage to obtain high-yield recombinant proteins (Figs. 2A&2B), and then performed circular dichroism spectroscopy experiments. CD spectra analysis confirmed predictions of the helical domain and we purified the helical domain of CHR with high yield. We could not crystalize the region which may need more length optimization. Then, we showed that the domain of CHR mainly composed of this helical domain associates with the C-terminus of BMAL1 and represses transcriptional activity of B/C complexes. And the N-terminus could help CHR located in the nuclei without any known NLS sequence. Nonetheless, in the CHRN sequences, there are ten arginine and four lysine, which have a potential to compose a novel NLS. Although CHRC had no obvious effects on B/C complex activity, this flexible domain may stabilize the interaction between CHR and BMAL1 or may form complexes with other proteins.

Recently, biochemical analysis and cell-based functional assays have helped us understand how negative factors inhibit the B/C complex-dependent transcriptional activation. Duong and colleagues reported that PERs form large complexes with more than ten other proteins, including many known clock proteins BMAL1, CLOCK, CRY1/2,CK1δ/ε, and NONO and other proteins such as PSF, SIN3, and HDAC around the E-box region on chromatin (34). The large protein complex deacetylates histone 3 and 4, and then represses B/C complex transcription. In contrast, the molecular PER-PSF-SIN-HDAC axis antagonizes acetyl transferase activity of CLOCK, offering another way that PER2 functions as a repressor (16). Ye and colleagues found that binding of PERs to the BMAL1/CLOCK/CRY complex removed BMAL1 from the E-box containing promoter regions of ccgs, resulting in loss of the transcriptional activation. Thus, PER proteins could act both ways to repress transcription (Fig.7). Differences in PER protein activity may depend on specific proteins PERs interacting with, and more work is required to understand these differences.

**Figure 7.**
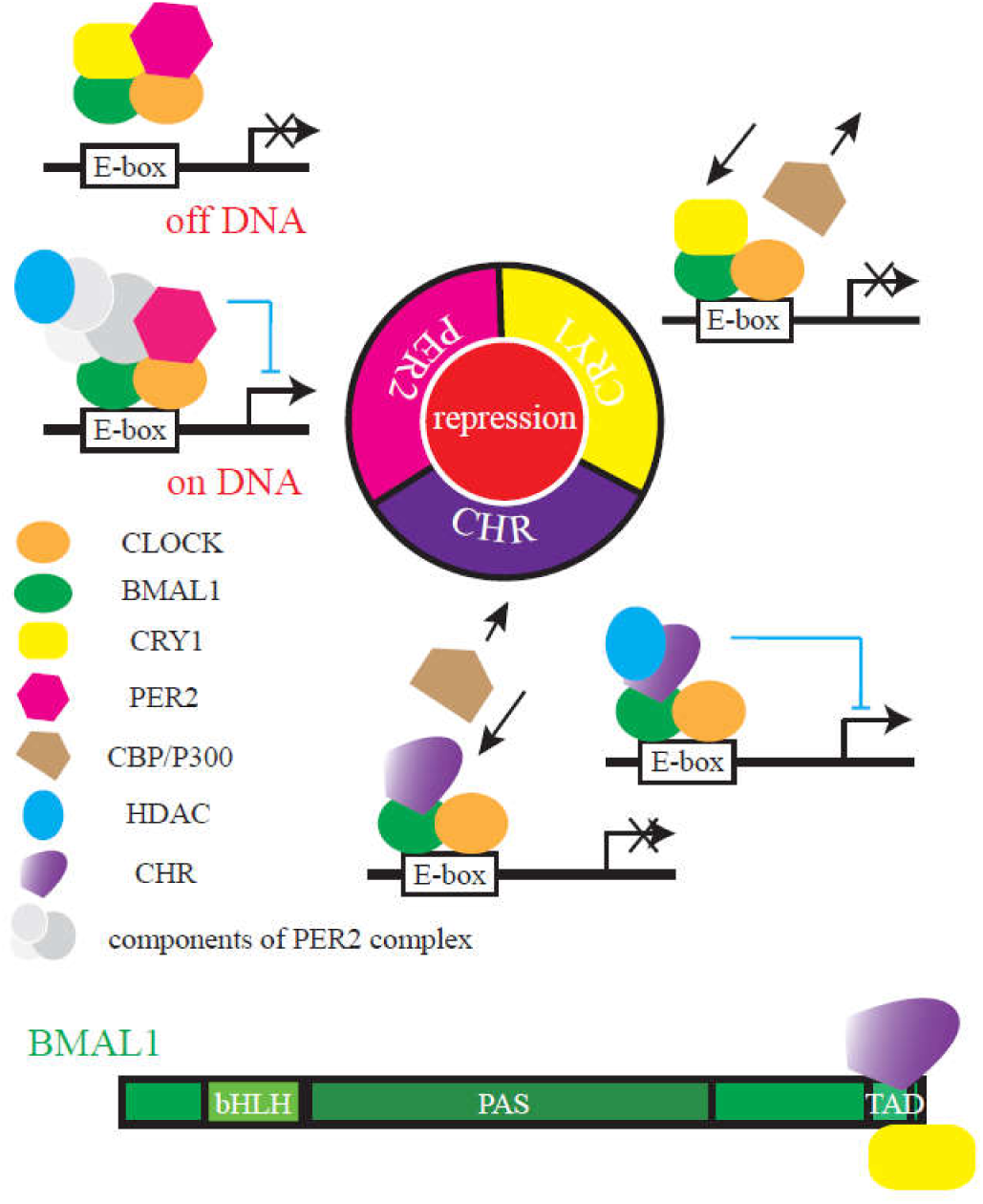
A molecular repression model of how CHR functions as a repressor in the mammalian circadian clock. During repression, PER2 can form large complexes to recruit HDAC or displace the B/C complex away from chromatin; CRY interacts with BMAL1 to restrict it from transcriptional co-activators; CHR binds to BMAL1 to either recruit HDAC or disrupt the recruitment of co-activators. PER proteins associate with the PAS domain of BMAL1; CRY1 and CHR bind to the TAD domain in the distal C-terminus of BMAL1. Under these different interacting modes, PER, CRY and CHR function differentially to repress the circadian clock.

CRYs function as a repressor independently of PER proteins. CRY1 is a stronger repressor than CRY2 during negative feedback (22), while CRY2 may repress transcription cooperatively with PER proteins. CRY1 competes with the transcriptional activator, CBP/p300 for binding to BMAL1 on chromatin (25). Early studies identified that the last 43 aa of BMAL1 (632 and 626 aa for mouse and human BMAL1, respectively) are required for association with CRY1 (23), and Xu and colleagues found that an L606A/L607A mutant at the BMAL1 C-terminus not only disrupted the association between CRY1 and BMAL1, but also abolished circadian cycling (25). The extreme end of BMAL1 is sequestered by CRY1 from transcriptional co-activators CBP/p300. Therefore, CRY1 proteins repress transcription differently than PER proteins.

Using co-IP, we reported that the 579-626aa region of hBMAL1 associates with the helical domain of CHR. Unlike previous reports that CHR binds to a different region of BMAL1 from where CRY proteins bind (14), our data supported that CHR and CRY associates to the same TAD domain of BMAL1 (Fig. 4).Goriki et al. reported that repression of CHR is HDAC-dependent (13). Collectively, we proposed a model by which PERs, CRYs, and CHR repress B/C transcriptional complexes (Fig. 7). In brief, PER can either form large complexes to recruit HDAC or displace the B/C complex away from chromatin; CRY and CHR interact with the extreme C-terminal end of BMAL1 to block it from transcriptional co-activators;CHR also recruits HDAC to disrupt the activation. PER proteins associate with the PAS domain of BMAL1 and CLOCK proteins (23), and CRY1 and CHR bind to the C-terminus of BMAL1(Fig. 7). Under such interacting modes, PER, CRY, and CHR differentially act as transcriptional repressors to regulate clock functions.

From the model we summarized above, it remains undetermined whether CHR can replace CRYs in the mammalian circadian clock. Studies from genetically modified animal models indicate abolished locomotor rhythms in *cry*1/*cry2* double knockout mice (35, 36). However, SCN slices from neonate *cry*1/*cry2* mice had robust circadian luminescence rhythm (37, 38). It is conceivable that CHR may compensate for loss of CRY proteins during early development, but later *ciart* expressed specific tissue. More work with CHR expression, especially the SCN in the *cry*1/*cry2* adult mice may provide important clues to solve this puzzle.

## Acknowledgments

This work was financially supported by grants from the National Natural Science Foundation of China (31771302 to XQ), the Anhui Provincial Natural Science Foundation (1608085MH212 to XQ). Ximing Qin acknowledges the start-up fund provided by Anhui University. We are also grateful to the staff for providing technical support with using the facility of Institutes of Physical Science and Information Technology.

